# Robustness increases heritability: implications for familial disease

**DOI:** 10.1101/2022.08.18.504431

**Authors:** Steven A. Frank

**Affiliations:** Department of Ecology and Evolutionary Biology, University of California, Irvine, CA 92697–2525, USA

**Keywords:** Quantitative genetics, genetic variability, tolerance, homeostasis, developmental variability

## Abstract

Robustness protects organisms in two ways. Homeostatic buffering lowers the variation of traits caused by internal or external perturbations. Tolerance reduces the consequences of bad situations, such as extreme phenotypes or infections. This article shows that both types of robustness increase the heritability of protected traits. Additionally, robustness strongly increases the heritability of disease. Perhaps the natural tendency for organisms to protect robustly against perturbations partly explains the high heritability that occurs for some diseases.

## Introduction

Robustness protects against perturbation. For example, checkpoints in the cell cycle sense certain types of DNA damage and prevent clonal expansion of aberrant cells. Robustness mechanisms reduce disease within individuals and, by protecting traits against the consequences of perturbation, may also influence patterns of genetic and developmental variability.^1^

In this article, I show that robustness alters variability in ways that increase heritability. I start with the heritability of any trait protected by robustness. I then describe how robustness enhances the heritability of disease.^2,3^

### Heritability

Consider a simple description of the heritability of a trait, *z*, as the ratio of the genetic variance, 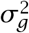, to the total trait variance, 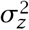, as

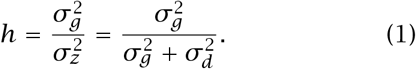

I split the total variance into a genetic component and a developmental component, 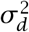. Nongenetic aspects are often called the *environmental* component. Here, I use *development* to emphasize processes within the individual that may be protected by robustness mechanisms, such as perturbations that alter the ways in which disease may develop within an individual.

### Increase in heritability

#### Robustness reduces developmental variation

Robustness mechanisms tend to buffer developmental fluctuations and random perturbations within individuals. For example, new DNA damage or perturbations to cellular environments can increase the tendency for clonal expansions. Cell cycle check-points act as homeostatic robustness mechanisms that reduce developmental variation in cell growth rates.

From eqn 1, any homeostatic robustness mechanism that reduces the developmental variance, 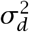, causes an increase in heritability when holding constant the genetic variance, 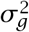. However, I show in a later section that a decrease in 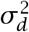 often decreases 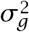. Evaluating the ultimate consequence of reducing developmental variability requires some analysis. It turns out that decreasing developmental variability does tend to increase heritability.

#### Robustness reduces deleterious effects

Robustness may raise heritability by reducing the effect of a potentially deleterious trait. For example, the heat shock protein Hsp90 helps amino acid sequences to fold properly into functional proteins. In the absence of Hsp90, a particular sequence may misfold. In the presence of Hsp90, the same sequence may fold properly.^4,5^

In general, as the deleterious effect of trait variants declines, the population will accumulate more genetic variation for that trait. For example, mutations leading to poorly folding amino acid sequences will be removed from the population relatively slowly when protected by the enhanced folding efficacy caused by Hsp90. Over time, the genetic variance, 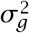, that causes poor folding in the absence of Hsp90 will increase.^4,5^

Any robustness mechanism that reduces the deleterious effects of trait variation will have the same consequence of raising 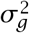. In eqn 1, as 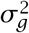 rises, heritability increases when holding all else constant.

However, robustness mechanisms that reduce deleterious effects alter the relation between character values and fitness. To understand the full consequences, we must analyze how all variables respond evolutionarily to changes in robustness. In a later section, I show that reducing deleterious effects and increasing an individual’s tolerance to character variation does typically raise heritability.

### Heritability of disease

This section highlights properties of key variables and definitions to evaluate the heritability of disease. The following sections show how robustness increases the heritability of disease. All models focus on asexual haploid genetics to emphasize the main qualitative trends in the simplest way.

#### Offspring phenotype given parental phenotype

Phenotype is the sum of independent genetic and developmental components, written as *z* = *g* + *d*. Suppose those components have normal distributions, 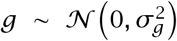 and 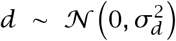, so that 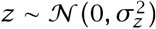, with 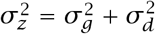.

To analyze the heritability of disease, we need to analyze progeny phenotype given parental phenotype. The first step is to describe the conditional distribution for the genetic value of the parent given its phenotype, *g*|*z*. From the derivation in the Supplementary Information,^6^ we obtain

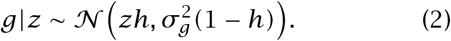

Progeny inherit their parent’s genotype, *g*|*z*. For a progeny’s phenotype given its parent’s phenotype, the value is *y*|*z* = *g*|*z* + *d*, in which progeny developmental component, *d*, is independent of its inherited genetic value. Thus, the conditional distribution of progeny phenotype, *y*, given parental phenotype, *z*, is

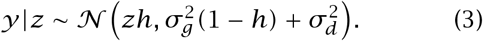

#### Heritability of disease

Suppose any phenotype greater than *z*^∗^ suffers disease. To simplify the analysis and without loss of generality, I limit disease to the upper tail of the phenotypic distribution. Let the familial measure of disease, *F*, be the probability that an offspring suffers disease, *y*|*z > z*^∗^, given that its parent suffered disease, *z > z*^∗^. Let the population-wide measure of disease, *D*, be the frequency of disease in the population, which is the probability that *z > z*^∗^. Define the heritability of disease as

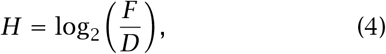

which is log_2_ of the odds ratio of offspring disease given parent disease, *F*, relative to the baseline disease probability in the population, *D*. Alternatively, one can use the odds ratio directly

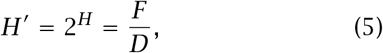

as in Fig. 1.

**Figure 1:**
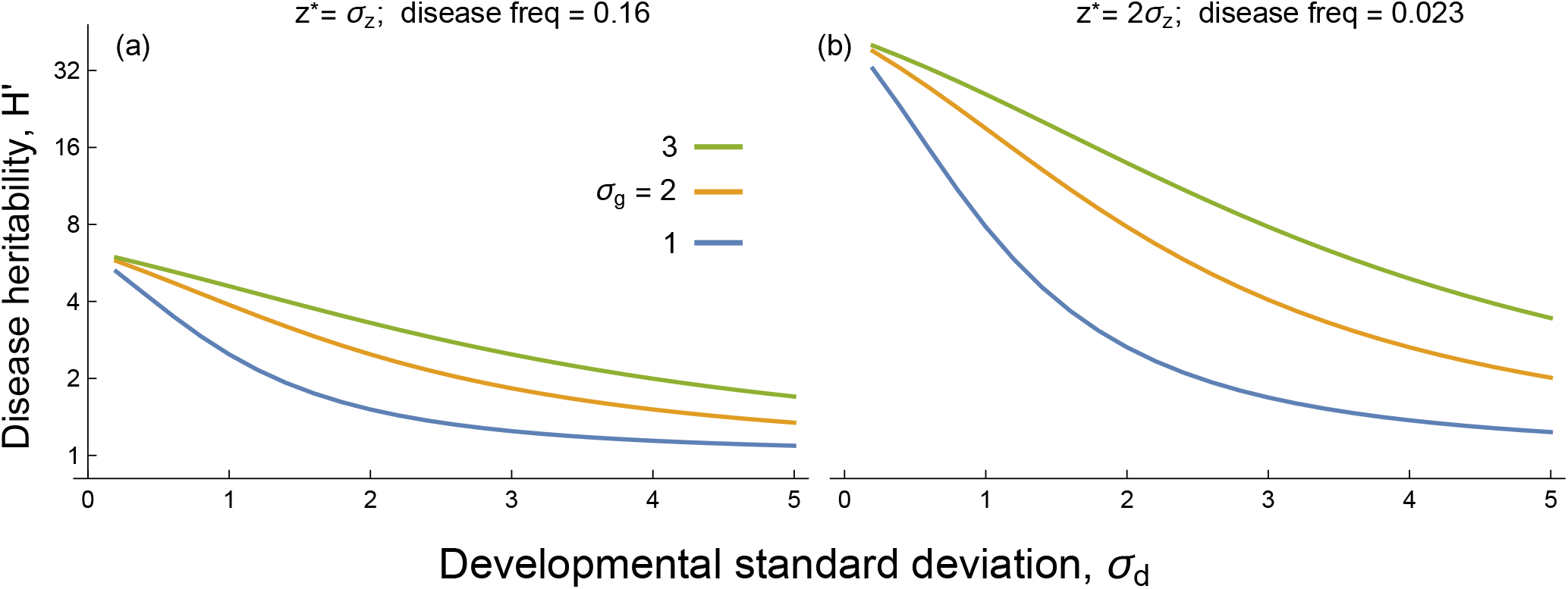
Robustness increases the heritability of disease. Typically, robustness tends to decrease developmental variance, 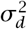, increase genetic variance, 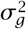, and decrease the frequency of disease, *D*. The curves in both panels show the heritability of disease, *H*^′^, as the odds ratio of *F*, the probability that an offspring suffers disease given that its parent suffered disease, relative to *D*, the population-wide probability of disease. In all cases, any decrease in developmental variance, 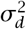, caused by robustness leads to an increase in the heritability of disease. The different colored curves in each panel show an increase in genetic variance from bottom to top. Any increase in genetic variance caused by robustness leads to an increase the heritability of disease. Panel (b) has a lower population-wide disease frequency than panel (a), because an increase in the phenotypic threshold for disease, *z*^∗^, associates with fewer individuals in the diseased upper tail of the phenotypic distribution. By setting the threshold to multiples of the phenotypic standard deviation, *σ*_*z*_, the disease frequency is constant within each panel over the various values of genetic and developmental variance.

The definition, simple in concept, requires five calculations. The Supplementary Information^6^ shows the derivation. Here, I emphasize the main steps. In the first step, the probability that *y*|*z > z*^∗^ is the upper tail above *z*^∗^ of the distribution in eqn 3. In the second step, we weight the probability that *y*|*z > z*^∗^ by the probability that the parent’s value is *z* and then sum over all parental values *z > z*^∗^. In the third step, we divide by the probability that *z > z*^∗^. These three steps give us *F*.

The fourth step sets the population-wide probability of disease, *D*, as the probability that *z > z*^∗^. The fifth step applies eqn 4 to obtain the log_2_ odds ratio of the probability that an offspring suffers disease given that its parent suffered disease, *F*, relative to the population-wide probability of disease, *D*.

### Robustness increases heritability of disease

The final value for the heritability of disease depends on the genetic and developmental variances, 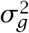 and 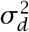, and on *z*^∗^ *>* 0, the threshold phenotypic value to be classified as disease.

Figure 1 shows how changes in one factor alter the heritability of disease when holding all other factors constant. Robustness may decrease developmental variance, increase genetic variance, or increase the phenotypic threshold for disease. The figure shows that any of these trends typically causes an increase in the heritability of disease when holding the other factors constant.

This partial analysis clarifies the components of the system. The following section considers how the various factors influence each other as they respond evolutionarily to changes in robustness.

### Evolutionary feedback

Full analysis must consider how changes in one aspect of robustness or variation alter other components. Consider, for example, a robustness mechanism that provides greater tolerance for extreme phenotypes. Weakened selection increases the genetic variance, increasing the frequency of individuals with phenotypes above a threshold, *z*^∗^. However, with greater tolerance, the threshold *z*^∗^ may no longer be the correct transition point to disease. Instead, the threshold shifts to a larger value.

How does robustness influence the heritability of disease when taking into account these feedbacks between the various key values? This section illustrates how to solve this problem. Each subsection considers a different robustness mechanism and its consequences.

#### Background

We need a few general definitions and results before turning to specific robustness mechanisms. To begin, define the fitness of a phenotype, *z*, as

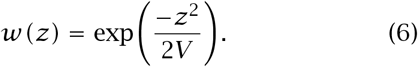

This expression assumes stabilizing selection with a maximum at *z* = 0. Then we can write the expected fitness of an individual with genetic value *g* as

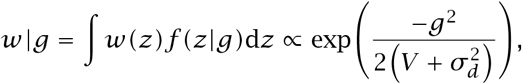

in which 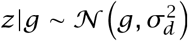 describes the developmental variation that determines the distribution of phenotype values, *z*, for a given genetic value, *g*, the function *f* is the probability density function, and ‘∝’ denotes *proportional to*.

To relate the genetic variance, 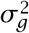, to the intensity of selection and the developmental variance, 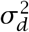, assume that the population is at mutation-selection balance. A classic model of mutation-selection equilibrium assumes that mutations causing a change in genetic value of ±*c* happen with frequency *µ* in each generation, and the selection intensity is set by the scaling of character value, *c*, and the denominator of the exponential that defines fitness.^7^–9 For example, using *w*|*g* for the fitness of genotypes, the selection intensity is 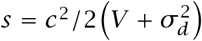.

When mutation is stronger than selection, 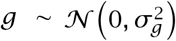, with

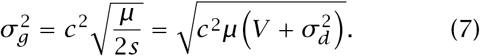

This expression relates the genetic variance, the developmental variance, and the intensity of selection on phenotypes given by *V*. The Supplementary Information^6^ provides details and also analyzes the case in which selection is stronger than mutation.^9,10^

Finally, we must consider how the threshold for disease, *z*^∗^, may change as evolution alters the other factors. I will assume that *z*^∗^ *>* 0 is the phenotypic value at which fitness is 0 *< γ <* 1, ignoring the symmetric point in the lower tail to keep things simple. Using *w(z)*, the value of *z* at which fitness is *γ* is

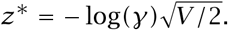

#### Increase in tolerance of phenotypic variants

In eqn 6, an increase in *V* flattens the fitness surface and weakens selection for a given phenotypic deviation from peak fitness. Weakened selection associated with larger *V* increases the genetic variance (eqn 7). A rise in genetic variance enhances heritability. Using the equilibrium genetic variance in eqn 7,

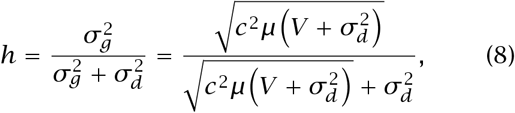

which increases with a rise in *V* (Fig. 2a).

**Figure 2:**
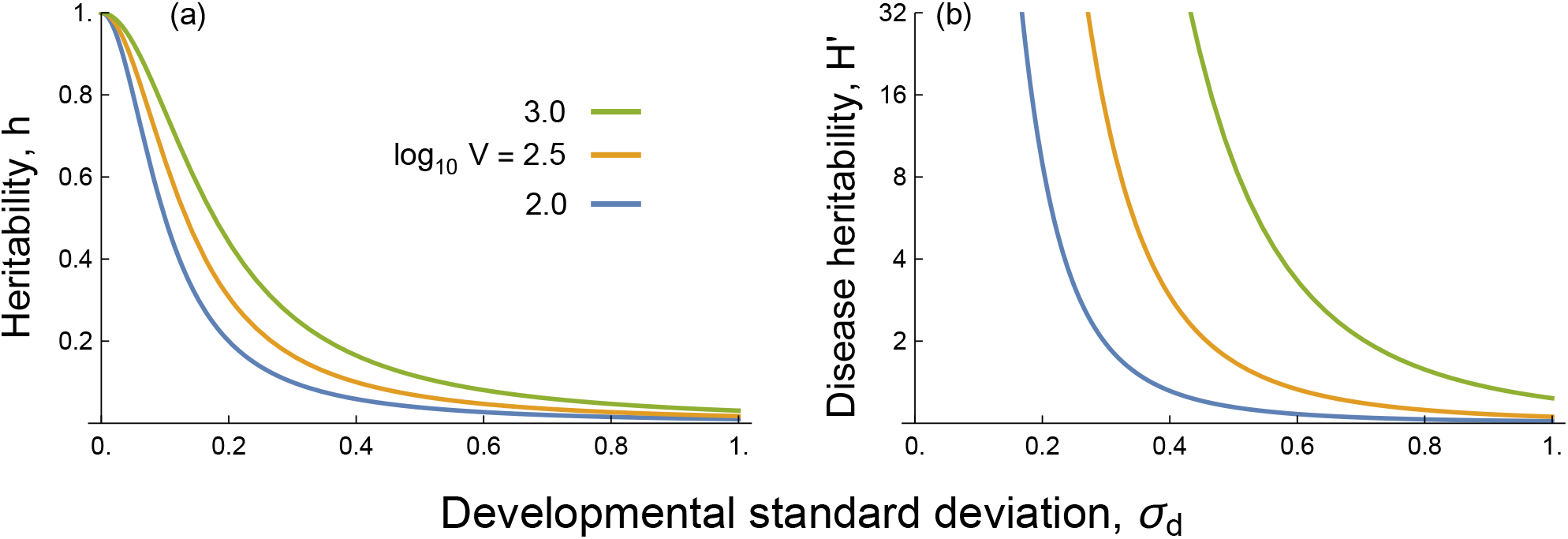
Evolutionary equilibrium accounting for interactions between variables. (a) The general heritability from eqn 8. (b) The heritability of disease, calculated by the steps outlined in the text and demonstrated in the Supplementary Information.^6^ Robustness causes decreasing developmental variance with decline in *σ*_*d*_ or weakening intensity of selection by tolerance with rising *V*. Either process increases heritability. For both panels, *c* = 0.1 and *µ* = 10^−4^. In (b), *γ* = 0.9.

#### Decrease in developmental variance

Heritability increases with a decline in developmental variance because 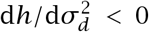, as shown in the Supplementary Information.^6^ The magnitude of the decline in heritability will depend on the parameters *c, µ*, and *V*. From eqn 8, we can plot heritability versus developmental variance for different values of *V*, as shown in Fig. 2a.

#### Heritability of disease

Figure 2b shows that increasing robustness raises the heritability of disease. In particular, a greater tolerance of phenotypic variation, increasing *V*, and a homeostatic decrease in developmental variance, declining 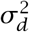, increase the heritability of disease. In Fig. 2b, the different component factors adjust to each other, leading to an evolutionary equilibrium, whereas in Fig. 1 each factor was changed holding the others constant. The same qualitative trends hold in the two cases.

## Discussion

A good test would compare two populations. The more robust population would have greater tolerance for phenotypic variants or greater homeostatic reduction of developmental variants. The theory predicts that the more robust population would have greater general heritability of the tolerated or variant trait and greater heritability of disease.

I do not know of any existing data that provide such comparison. I mention a few promising candidates for future study.

Plant tolerance to herbivory and disease has been widely discussed.^11^–13 Many cases of phenotypic plasticity are considered within the context of tolerating environmental challenges.^14^ Comparison between populations or species that differ in tolerance may be interesting.

Microbial exodigestion arises when individual cells secrete enzymes.^15^ When diffusion is sufficiently fast, local digestion depends on the average secretion rate per cell in the neighborhood, independently of the variation between cells. The developmental variation of secretion rate is robustly buffered by local averaging. By contrast, in viscous environments with slow diffusion, the effective neighborhood shrinks. Group averaging no longer buffers individual cellular variation.^16^

Comparing fast versus slow diffusion, the fast diffusion situation favors greater heritability of secretion rate per cell and greater heritability of extreme phenotypic variants that may suffer reduced fitness, an expression of disease.

Protections against cancer seem to differ between species.^17^ Larger or longer-lived species may have greater protection for a given tissue. If so, then the heritability of disease may be greater in those species with greater protection. However, because the genetics of disease likely differs between such species, clarity about mechanism may be required to make a clear comparison.

## Supplementary information

Two files are available online.^6^ The file heritable.nb provides a Wolfram Mathematica notebook that can be used to follow the steps in derivations and to evaluate alternative quantitative assumptions. A printed version of that notebook in the file heritable.pdf can be read without using the Mathematica software.^18^

